# Males perceive honest information from female released sex pheromone in a moth

**DOI:** 10.1101/2021.04.14.439460

**Authors:** Golov Yiftach, Liberzon Alexander, Gurka Roi, Soroker Victoria, Jurenka Russell, R Harari Ally

## Abstract

There is accumulating evidence that male insects advertise their quality to conspecific females through pheromones. However, most studies of female released sex pheromone assume information transfer regarding merely the species of the female and her mating status. We show that more and precise information is conveyed through the female sex pheromone, positioning it as an honest sexual trait. We demonstrate that females in bad physical conditions (small, starved or old) lay significantly fewer eggs than females in good conditions (large, fed or young). The ratio of the sex pheromone blend in gland extracts of female pink bollworm moths accurately describes the female phenotypic condition whereas the pheromone amount in the glands fails to provide an honest signal of quality. Moreover, males use the female released pheromone blend to choose their mates and approach females that signal higher reproductive potential. In addition, surrogating the female effect, using synthetic pheromone blend that represents that of higher quality females (0.6:0.4 ZZ:ZE) more males were attracted to this blend than to the blend representing the population mean (0.5:0.5 ZZ:ZE). Both, female advertisement for males and the male choosiness, suggest that pheromones have evolved as sexual traits under directional, sexual selection.

## INTRODUCTION

Chemical communication via pheromones is commonly used across taxa (Wyatt 2014). Whereas most previous studies considered the significance of pheromone mainly in species-(Löfstedt et al. 1991; Baker 2002; Mas and Jallon 2005) and sex-recognition (Löfstedt et al. 1990; Svensson 1996; Johansson and Jones 2007; Bacquet et al. 2015), more recent studies provide evidence that sex pheromones convey honest signals on the individual’s condition and its quality as a mate (Johansson et al. 2005; Harari et al. 2009; Foster and Johnson 2011; Harari et al. 2011; Harari and Steinitz 2013; Chemnitz et al. 2015).

Pheromones, acting for species or sex recognition may be under stabilizing selection (Collins and Cardé 1985; Löfstedt et al. 1993; Phelan 1997), therefore are expected to show little to no individual variation to restrict the high cost of possible identification mistakes. However, when signaling mate quality, pheromones are expected to vary between individuals of the same species and reflect differences in phenotypic conditions, conveying the accurate alteration in the quality of the sender (Allison and Cardé 2008; Harari et al. 2011; Ruther et al. 2009). Furthermore, as the sender of the pheromone will benefit greatly from the signal, these signals are subject to cheating and therefore are likely to be costly (Zahavi 1975, 1977). In general, costly traits are condition dependent, such that individuals with better phenotypic conditions are able to produce signals that attract more mates (Grafen 1990; Johnstone 1995).

Indeed, with the understanding that costly male sexual traits have coevolved with the female preference for elaborated signals (Rowe and Houle1996; Kotiaho 2000; see review in Cotton et al. 2004), the positive correlation between male’s sexual trait (pheromone included) and the male’s reproductive success has been well established (Rantala et al. 2003; Fisher and Rosenthal 2006; Blaul and Ruther 2011; Nieberding et al. 2012; Pölkki et al. 2012; South et al. 2011; Chemnitz et al. 2015).

However, studies supporting similar correlation between female produced pheromones and their reproductive success are extremely scarce (Jaffe et al. 2007; Harari et al. 2011). Possible explanations for this lack are (1) the general assumption of sexual selection theory that females are the choosy sex as they are limited in gametes, whereas males are limited in mating opportunities (Trivers 1972; Bateman 1948). (2) The cost of pheromone produced by female moths is traditionally assumed to be negligible (Greenfield 1981; Cardé and Baker 1984; Svensson 1996; Allison and Cardé 2008; Foster and Johnson 2011; Kokko and Wong 2007), due to its specificity and high effectivity at minute amounts (Wyatt 2014). (3) The conception that stabilizing selection is operated on female emitted pheromones, thus, only little variance is expected (van Wijk et al. 2017). As such, female moths’ pheromones have been taken as incapable to honestly signal the females’ phenotypic conditions and may serve solely to direct mate searching males to intraspecific females. Nevertheless, male choice of females is common in many taxa (Bonduriansky 2001), and is expected to be the norm in moths as males are sperm limited (Friedländer et al. 2005; Teng and Zhang 2009). Yet, studies on male preference for females of better condition based on their pheromone characteristics are extremely limited.

Intraspecific variation in female moths’ sex pheromones has been documented (Groot et al. 2014; Löfstedt et al. 1986;) and more so with the advanced technology that allows to quantify the pheromone amount and ratio in an individual female (Levi-Zada et al. 2020). Moreover, sex pheromones are subjected to changes in quantity and quality during the females’ lifespan (Aliniazee and Stafford 1971; Levi-Zada et al. 2020; Liu and Haynes 1994; Webster and Cardé 1982) and there is evidence suggesting that female sex pheromones are affected by the female’s physical condition (Foster and Johnson 2011; Harari et al. 2011). All the above mentioned suggest a cost of pheromone production in female moths, proposing female sex pheromones as honest sexual signals that may be evaluated by mate searching males. Thus establishing sex pheromones of female moths as a sexual trait affected by sexual selection pressures.

Here we tested if pheromones, produced by female moths, honestly reflect the female’s condition and whether this information is used by males to choose females in better conditions. Three condition-dependent traits, known to affect female fecundity, were chosen: size (Harari et al. 2003; Honěk 1993; Li et al. 2005), age at mating (Torres-Vila et al. 2002), and diet (Leahy and Andow 1994; Wäckers et al. 2007). We used the pink bollworm (*Pectinophora gossypiella*) as a model species as the female pheromone blend is known and constructed from two isomer components, both in detectable amounts using gas chromatography (Hummel et al. 1973).

## Methods

*Pectinophora gossypiella* larvae were reared in climate rooms at 27±1° C, 14:10 L:D, on an artificial diet (Stonefly Heliothis diet, Ward’s Science). Males and females were sexed during their last larval stage by the detection of a black line on the 6^th^ abdominal segment, representing the developing testicles. Males and females were housed separately and newly-emerged moths were removed daily and placed into age cohort single sex cages. Male moths were fed on ∼10% sucrose solution provided ad libitum.

### Females’ phenotypic conditions

Three distinct physical conditions were chosen: Size, age and diet. (1) Size of females was determined during the pupal stage. Females divided into size groups of either small (range: 0.005-0.008, mean: 0.007 g), medium (range: 0.009-0.014, mean: 0.012) or large (range: 0.015-0.030, mean: 0.023 g). Only large and small females were used in this study. (2) Age of females was measured starting from the day of their emerging as adults. Three-days-old females were defined as young females, whereas, seven-days-old females were defined as old females. (3) Diet of females was manipulated by providing either 15% sugar solution to “fed” females or water only to “starved” females. Both, sugar solution and water were provided to females ad libitum, starting from the day of their emerging as adults..

Based on size, age and diets, females were divided into four condition-dependent categories: the “Size” category includes large and small females that are three days old and sugar fed. The “Age” category includes young and old females that are large and sugar fed. The “Diet” category includes fed and starved females that are large and three days old. “Diet and small” category, includes fed and starved females that are small and young. Note that the female phenotypic-condition: large, young and fed takes part in the categories “size”, “age” and “diet”. The phenotype small, old and starved was not included in the analyses, as most small and starved females did not survive for old days (7 days).

### Condition-dependent fecundity

In order to verify the effect of the female phenotypic-condition on her fecundity, we counted the eggs laid by each female in her lifetime, in all phenotypic conditions (Table 1).

**Table 1.** The female phenotypic conditions used to measure condition-dependent fecundity.

| Treatment of Phenotypic-condition | Size |  | Age |  | Diet |  |
| --- | --- | --- | --- | --- | --- | --- |
| Phenotypical types | Large | Small | Young | Old | Fed | Starved |
| Large, young and fed | X |  | X |  | X |  |
| Large, young and starved | X |  | X |  |  | X |
| Large, old and fed | X |  |  | X | X |  |
| Small, young and fed |  | X | X |  | X |  |
| Small, young and starved |  | X | X |  |  | X |
| Small, old and fed |  | X |  | X | X |  |

Upon emergence, 50 females of each phenotype (Table 1) were placed with 70 randomly selected males of the medium size. Males and females were used a day following their emergence. The pair, in copula, was carefully placed in a glass tube. Couples that stop mating within less than 30 minutes (time to transfer a full spermatophore) were discarded. Three hours later each couple was introduced to a plastic cup (150 cc), covered by a net. On top of the net, a towel paper was placed for oviposition. Sugar solution for fed females and water for starved females was provided through a glass tube sealed with cotton week, inserted through a hole in the cup wall. After the death of the female all the eggs deposited on the towel paper were counted.

### Pheromone extracts

In the pink bollworm a good correlation in ratio of components was found between glands extracted pheromone and the pheromone emitted by calling females (Allison and Cardé 2016). Thus, in order to verify the effect of the female phenotypic-condition on her pheromone characteristics we measure the amount and ratio of the two components in gland extracts of the different females (Table 2). Due to available sample size and technical limitation using GC, the pheromone characteristics in gland extracts were compared in pairs (two treatments of the same category at a time), according to the Y-tube dyad experiments.

**Table 2.** Testing males’ preference of females that vary in their phenotypic conditions u a Y-tube bioassay. Males preference of small and old females was not measure as small females rarely survive to old age (seven days).

| Treatments | Y- tube arm | Size |  | Age |  | Diet |  |
| --- | --- | --- | --- | --- | --- | --- | --- |
| Sub-groups |  | Large | Small | Young | Old | Fed | Starved |
| 1 | 1 | X |  | X |  | X |  |
|  | 2 |  | X | X |  | X |  |
| 2 | 1 | X |  | X |  | X |  |
|  | 2 | X |  |  | X | X |  |
| 3 | 1 | X |  | X |  | X |  |
|  | 2 | X |  | X |  |  | X |
| 4 | 1 |  | X | X |  | X |  |
|  | 2 |  | X | X |  |  | X |

Pheromone glands of females were extracted to quantitatively assess the pheromone’s absolute amount and the ratio of its components. Pheromone glands of virgin *P. gossypiella* females, representing the different treatments, were removed 2 hours after the start of the scotophase, extracted for 20 min in hexane with 25 ng tridecyl acetate as an internal standard. Gland extracts were analyzed with a Hewlett Packard 5890 GC coupled to a 5972 mass selective detector using a DB Wax capillary column (J & W Scientific, 30 m x 0.25 mm). The GC was programmed at 60°C for one minute, subsequently ramping at 5°C/min to the final temperature of 230°C which was held for 15 minutes. The mass spectrometer was set in single ion monitor mode for ions 43, 55, 67 and 81 (the 4 most abundant ions of Z,Z- and Z,E-7,11-16:OAc) and 43, 55, 61, and 69 (the 4 most abundant ions of tridecyl acetate).

### Male mate preference

#### 1. Calling females

In order to test whether male preferences for females is related to the female’s phenotypic-condition, we let randomly selected, sugar fed, three days old males to choose, in a Y-tube bio-assay, between females of two phenotypic-conditions (Table 2).

A Y-tube olfactometer (Bine, Weitzman Institute for Science, Rehovot, Israel) was used to test the attraction of the odor of virgin females that are varied in their phenotypic-condition. The olfactometer consisted of a central tube (20 cm long, 2.5 cm diameter) and two lateral arms (72°, 14 cm long, 2.5 cm diameter), each connected to a glass jar (250cc) containing 5 females via Teflon tubing (20 cm long, 0.5 cm diameter). A sieve was inlayed in the connecting glass tube, between the Teflon and glass tubing to prevent the female insects to escape. Charcoal purified and humidified air was passed into the olfactometer through a Teflon connection at 30 mL/sec. To avoid visual distraction for the moths, the jars containing the females were hidden from the males in the Y-tube. The male moth walked up to the two arm’s junction and its choice of arm was recorded when reached a predefined line, 12 cm from the junction. Males that did not choose within 3 min after releasing were removed and recorded. Between every five runs, the olfactometer arms were flipped (180°) to minimize positional effect. Tests were conducted in a dark room, 26°C, 2-4 hrs. into the scotophase, which was verified in preliminary observation as the time of courtship and mating. Females and males were transferred to the experimental room during the photophase to experience the onset of the scotophase in the experimental room, on time, allowing for 2 hrs. of acclimation time.

Generally, females call intermittently and both timing and duration of calling vary among females in the population (Harari et al. 2011). Therefore, to increase the chances of female calling throughout the experiment, five pink bollworm females were used as a source of pheromone in each arm of the olfactometer. Males preference for females in each phenotypic condition was repeated in three days with ∼30 males in each day. At the end of each day, the olfactometer parts were rinsed in soapy water and dried with ethanol (70%).

#### 2. Synthetic pheromone

In the olfactometer (as above) we tested the male preference for the synthetic pheromone components Z,Z-7,11-16:OAc (ZZ) and Z,E-7,11-16:OAc (ZE) (Pherobank, The Netherlands, Cat. Nos. 10523 and 19424) at ratio of 0.6:0.4 representing that of a high quality female vs. 0.5:0.5 ZZ:ZE, representing the pheromone profile of the population mean. The total concentration of the pheromone was 4 ng per µl of hexane. The 10 µl volume of the solution (equivalent to 5-7 females) with the desired ratios of components was spread on a 5×5 <u>cm</u> parafilm sheet that was used as a dispenser and kept in -18 c until used.

A sheet of each treatment served as volatile source in the olfactometer, one at each side (arm). Males were released, one at a time, down the wind at the long arm of the Y tube. Only males that reached the far end of one of the Y arms within 2 min were recorded. The direction of the odor source was rotated (left-right) every 5 runs to avoid positional bias. The paraphilm sheets containing the pheromones were changed every 30 - 40 minutes to ensure a continuous supply of pheromones blends.

### Statistical analyses

#### Condition-dependent fecundity

Preceding analysis of Mahalanobis Distances was conducted for each of the six phenotypic-condition variables (Table 1), in order to identify and exclude outliers before performing the comparison analysis. The data did not meet the assumptions of normality (p<0.05 in both, Levene’s Test for Homogeneity of Variances of multiple samples, and Shapiro–Wilk tests). Therefore, a two-sided nonparametric Kruskal– Wallis H test (α=0.05), followed by the post-hoc method of Steel–Dwass test were performed to compare the number of oviposited eggs among females that vary in their physical conditions (Table 1).

#### Pheromone characteristics

GC chemical analyses of the pheromone glands were conducted in dyads, in accord with the Y-tube behavioral assays. Full pairwise comparison of gland extracts among the different dyads (8 groups in total) is statistically not valid. Hence, the GC results were statistically conducted only on the same treatment, resulting in four comparisons (as in Table 2). This methodology allows to compare the two traits of the pheromone in the gland extracts: (i) the amount (ng) and (ii) the ratio of the two components (Z,Z/ Z,E -7,11-hexadecadienyl acetate). For the amount of the pheromone the assumption of homogeneity of variances was violated (Levene’s test, p<0.05), thus, the two-sided Mann-Whitney U Test was used (α=0.05). The ratio between the two pheromone components, (Z,Z) / (Z,E) was compared using a two-sided T-test (α=0.05), as all treatments meet the assumption of homogeneity of variances (Levene’s test, p>0.05).

The relationship between pheromone amount and its ratio of components. To test for a dependency between both traits of the pheromone (amount and ZZ/ZE ratio) at the individual level, a correlation (Spearman’s rank correlation coefficient) and a linear relationship (linear regression), analyses were performed for each of the four pairs (as in Table 2). The correlation was tested after excluding extreme outliers, using Mahalanobis Distances analysis.

##### Male preference

Contingency tests, after testing for homogeneity of variance (Chi-Square Test, p>0.05), was used to analyze the male preference to females in the olfactometer. The analysis included two types of comparisons: (i) within a group (same dyad treatment) for each of the four dyads, using a two-sided G^2^-test goodness of fit (DF=1). (ii) Among all four dyads, using two-sided Fisher exact test for multiple comparison (DF=3), following by post-hoc comparison using two-sided Fisher exact test of 2×2 (DF=1), after Holm-Bonferroni adjustment.

## Results

### Condition-dependent fecundity

Lifetime fecundity of pink bollworm females is definitely condition dependent (Kruskal-Wallis test, H=111.17, p<0.0001, n=177) (Figure 1). Females in a very good physical condition (large, young and fed) had significantly more eggs than females that experienced reduced physical conditions. Deteriorating conditions in size, age or diet resulted in significant reduced fecundity of females. Among all physical conditions, size had a major effect on the number of eggs laid, and within size groups, starved females experienced the least lifetime fecundity (Large, young and fed - 103.2 ± 9.43 eggs, n=26; Large, old and fed - 65.87 ± 7.01 eggs, n= 31; Large, young and starved - 27.03 ±3.69 eggs, excluding one outlier, n=29 out of 30; Small, young and fed - 22.63 ± 3.0 eggs, excluding one outlier, n=30 out of 31; Small, old and fed - 10.29 ±1.0 eggs, n=31; Small, young and starved - 6.57 ± 0.89 eggs, n=30).

**Figure 1.**
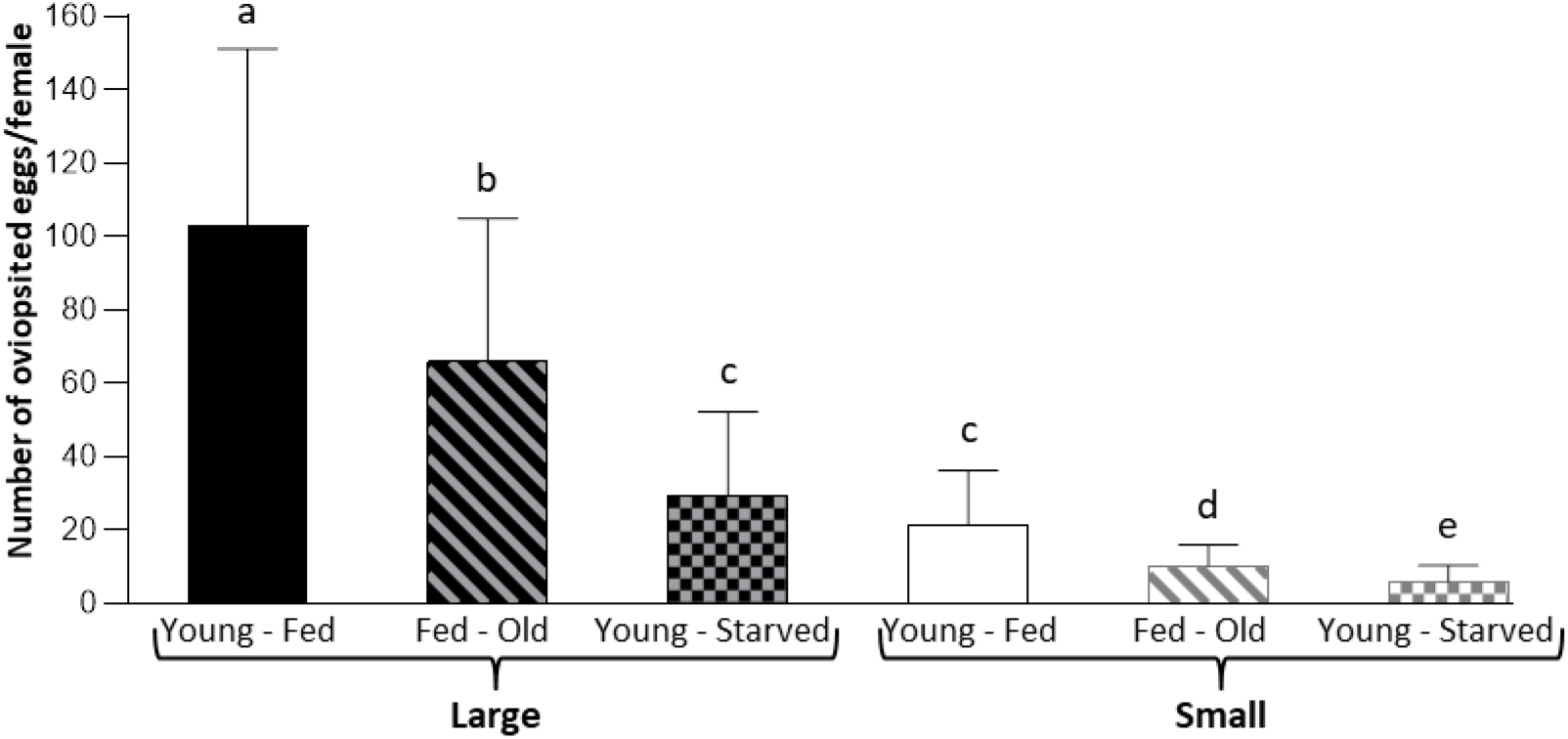
Condition dependent fecundity. Life time fecundity of 3 day old small (white columns) and large (black columns) pink bollworm females that fed on sugar solution (fed), on water (starved, displayed by diagonal strips) and old (7 days old, sugar fed, displayed by squared net) females 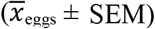. Kruskal–Wallis H test followed by Steel–Dwass Multiple Comparison test. Different letters indicate significant difference at p<0.05.

### Pheromone extracts

Comparisons of the pheromone characteristics were conducted within dyads (Figure 2). No significant differences were reveled in the amount of pheromone in three of the dyads tested. (Mann-Whitney U test: Mean ± SE, (1) Size - large, young and fed females: 9.58+2.24 ng, n=14 vs. Small, young and fed females: 6.56+1.21 ng, n=10; U=52.0; (2) Age - young, large and fed females: 8.16+0.81 ng, n=27 vs. old, large and fed females: 8.24 ± 1.84 ng, excluding two outliers, n=28 out of 30; U=359.5, p>0.05; (3) Diet – fed, large and young females: 12.78 ± 1.33 ng, n=14 vs. starved large and young females: 13.14 ± 1.16 ng, excluding one outlier, n=16 out of 17; U=107.5, p>0.05). However, a significant difference was found comparing the amount of pheromone in gland extracts of small, young females that differed in their diet (Fed, small and young females: 9.43 + 1.68 ng, excluding three outliers, n=21 out of 24 vs. Starved, small and young females: 2.11 ± 0.45 ng, n=11; U=28, p<0.0001). An analysis of the ratio between the two components of the pheromone blend, ZZ and ZE was conducted for each of the four dyads (Figure 3). The analysis revealed a significant difference between treatments in three among the four dyads: (1) Size-Large, young and fed females had significantly more of the ZZ component in their pheromone than small young and fed females (Levene’s Test: p>0.05, T-test: t=2.18, p=0.04; Mean ± SE of ZZ/ZE ratio, 0.535+0.001, n=14 and 0.504+0.012, n=10, respectively). (2) Age – Young, large and fed females had significantly more of the ZZ component in their pheromone than old, large and fed females (Levene’s Test: p>0.05, T-test: t=3.03, p<0.01; 0.494 ± 0.008, n=27 and 0.459 ± 0.008, n=30, respectively). (3) Diet – Fed, small and young females had significantly more of the ZZ component in their pheromone than starved, small and young females (Levene’s Test: p>0.05, T-test: 3.42, p<0.01; 0.546 ± 0.006, n=24 and 0.506 ± 0.012, n=11, respectively). However, no significant difference was detected in the ZZ/ZE ratio when large and young females differed in their diet (sugar solution vs water) (Levene’s Test: p>0.05, T-test: t=0.84, p>0.05; Fed, large and young females - 0.537 ± 0.014, n=14 and starved females - 0.535 ± 0.013, n=17).

**Figure 2.**
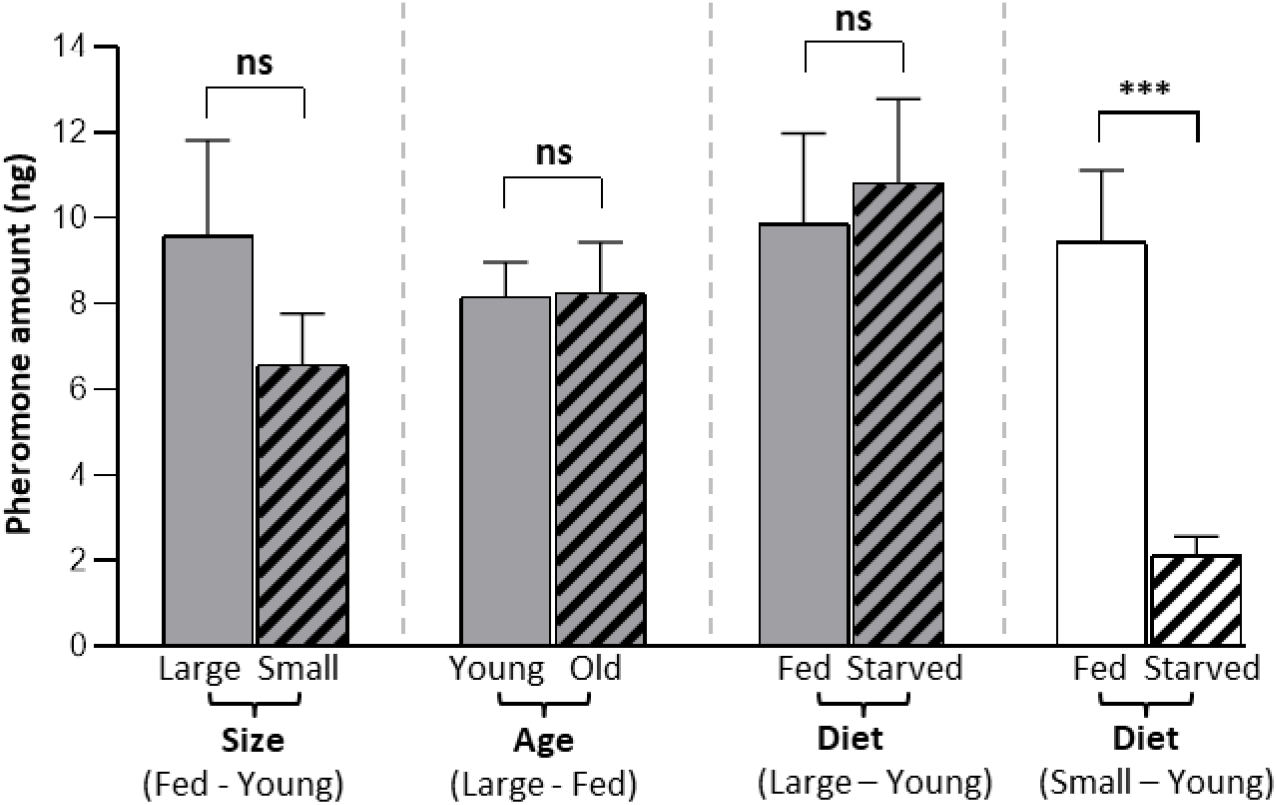
Pheromone amounts (ZZ + ZE) in pheromone glands of pink bollworm female taken from the indicated groups as described in methods. X-axis – the four dyads of treatments: (1) Size difference, (2) Age difference, (3) ‘Diet difference of large’ (grey columns) and (4) ‘Diet difference of small (white columns)’. The two sub-groups of each treatment displayed by clear and diagonal lines. *** indicates statistical significant difference between mutual sub-groups, Mann–whitney U test p<0.0001.

**Figure 3.**
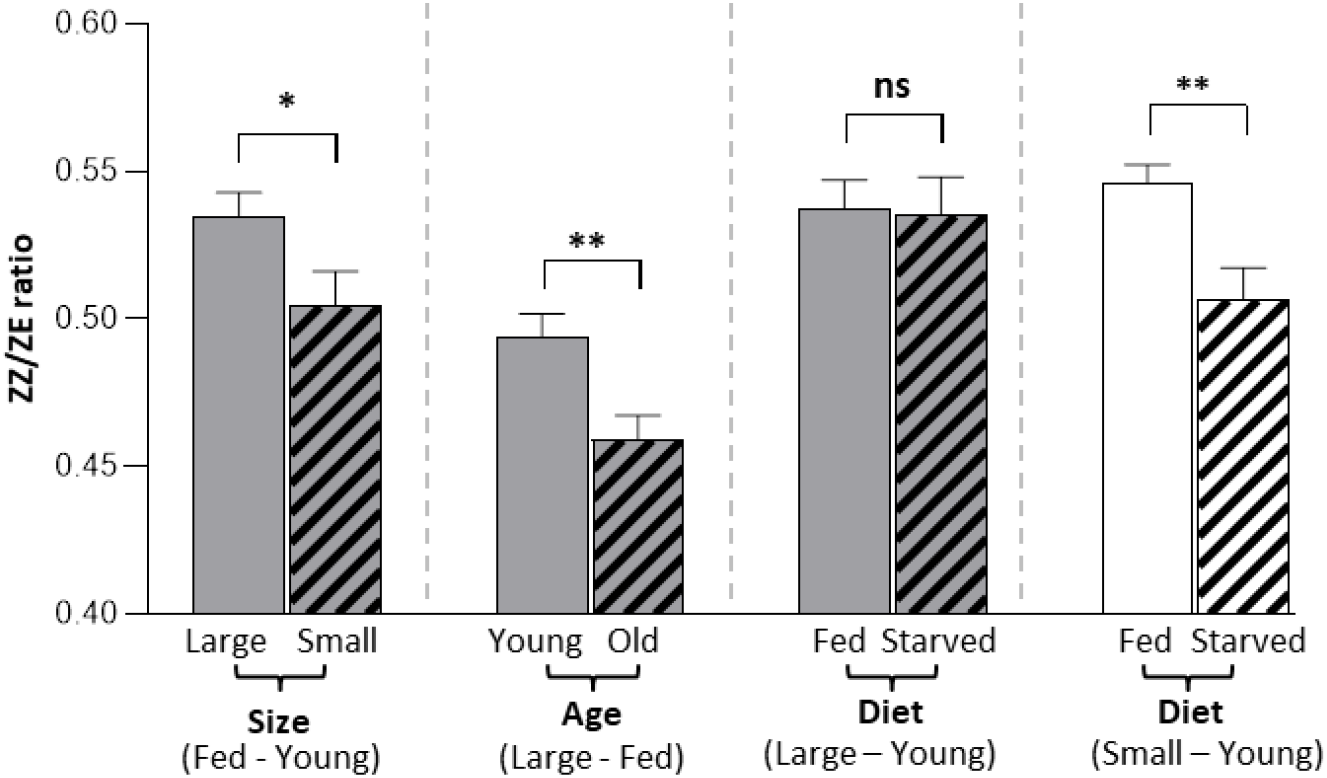
Ratio of pheromone components (ZZ/ZE ratio) found in the pheromone glands of pink bollworm taken from the indicated female groups as described in methods. Asterisks indicate statistical significance of difference between mutual sub-groups with * p < 0.05, ** p < 0.01, NS = not significant, T-test.

The relationship between pheromone amount and its ratio of components. Testing for a dependency between the amount of the pheromone and its ratio of components (Figure 4) revealed a significant positive correlation in all four dyads (Spearman’s rank correlation coefficient, p<0.05, after excluding one outlier in the size category, two outliers from the age category, and two outliers form large and diet category). Interestingly, in three out of the four dyads (Size, Age and Diet of small females, Figure 4.A, B, D, respectively), the amount and ratio of components, of the better quality females in each dyad (larger, younger and fed) area above the correlation line. This indicates a higher amount of pheromone and more of the ZZ component in high quality females than in inferior females, in these dyads (smaller, older and starved). Only in one dyad (fed and starved, young, large females, Figure 4.C) this tendency was not detected. In addition to the independent quantitative analyses (fecundity or pheromone traits), the integration of all parameters was explored using the multivariate descriptive technique of multiple correspondence analysis (MCA, see Appendix. 1 &2). First, at the treatment level, ‘size difference’ and ‘diet difference of small’ were the most distinctive groups (located farther from the point of intersection of coordinate axes, 0.0, 0.0). Moreover, the physiological effect of the two treatments, ‘age differencex2019; and ‘diet difference of large’ was relatively similar, as shown by the mutual proximity of these two groups. Yet the proximity of these two treatments to the feature of higher quality (mark by ‘+’) was different as the ‘diet difference’ was the closest treatment. Secondly, at the dyads level, the most distant groups were both of the ‘diet of small’.

**Figure 4.**
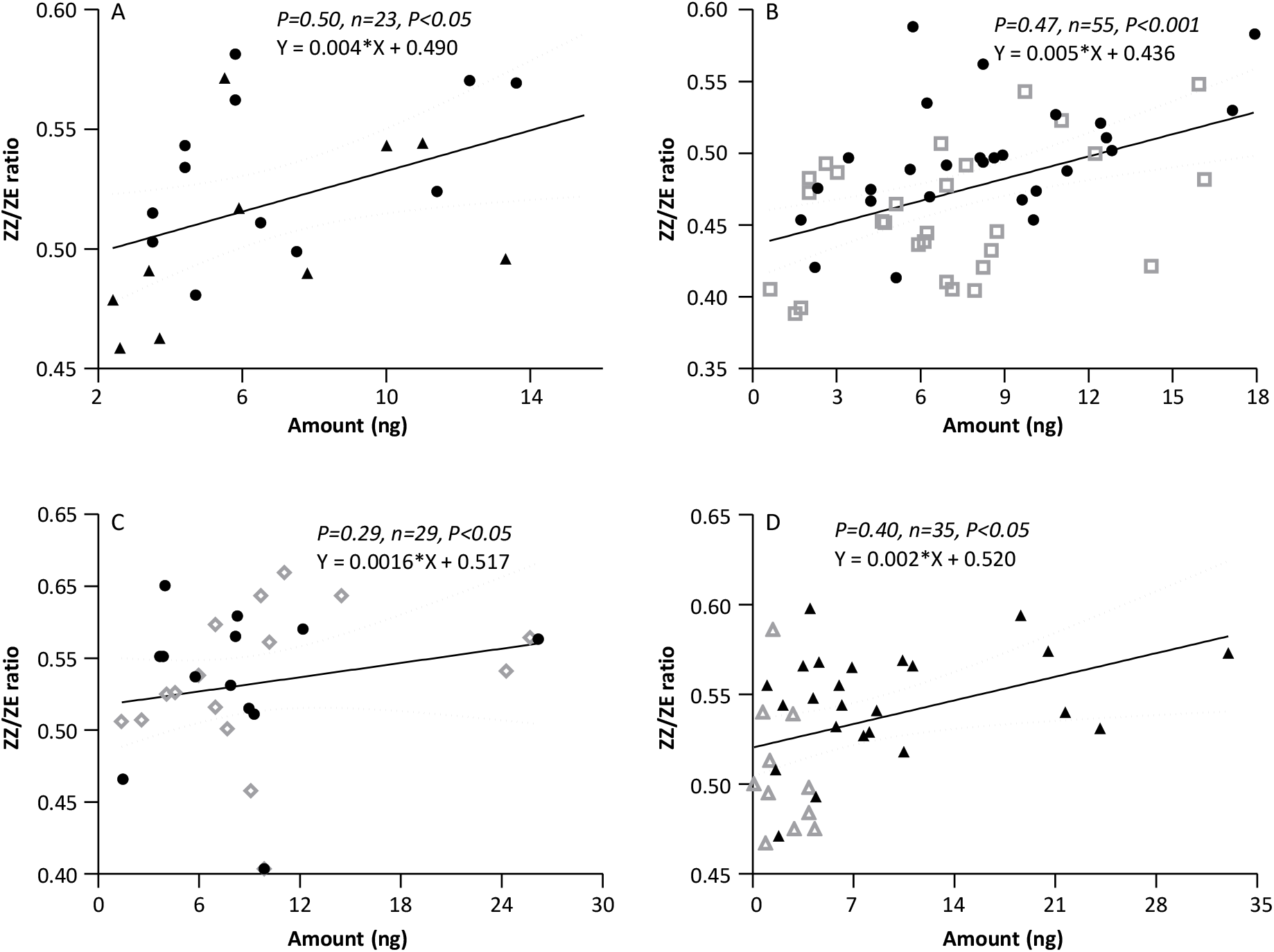
Relationship between the two traits of the pheromone gland extracts. X-axis: the amount (ng) of the extract pheromone (ZZ + ZE). Y-axis: the ratio between the two components, ZZ and ZE (ZZ/ZE ratio). A, all females of the ‘*Size*’ treatment, B, all females of the ‘*Age*’ treatment, C, all large females of the ‘*Diet*’ treatment, D, all small females of the ‘*Diet*’ treatment. Symbols represent the eight groups of females’ conditions as follow: large – young – fed (black circles), small – young – fed (black triangulars), large – old – fed (empty squares), large – young – starved (empty rhombus), and small – young – starved (empty triangulars). Spearman’s rank correlation coefficient test (α=5%). Solid lines display the liner regression model between the two variables; dashed line represents the 95% confidence interval (CI) of the linear model.

### Male mate preference

#### Live females

Males’ preference was tested in olfactometer assay of choice experiments of the four dyads (Figure 5). Testing the effect of the female size, a full factorial analysis between the females’ physical category conditions (x_1_) and the males’ preference (y_1_) revealed that each of the physical phenotypic categories condition had significantly different effect on males preference (Fisher exact test for multiple comparison, DF=3, n=348, p<0.05). The number of males who reached the arm leading to large females was significantly higher than to the arm leading to small females (71 vs 14 males, respectively, G-test p<0.0001, G^2^=40.75, p<0.0001; 14 males (14.3%) did not show a preference during the first 3 min of the experiment). Similarly, when testing the effect of female age on male preference, more males preferred young females over old ones (53 vs 25 males, respectively, G-test p<0.001, G^2^=10.28, p<0.01; 20 males (20.4%) did not show a preference during the first 3 min of the experiment). Testing the effect of diet on male preference on both large and small females, revealed that more males preferred the fed females over starved ones (Large and young females: 58 vs 29 males, respectively, G-test p<0.01, G^2^=9.85, p<0.001; 20 males (18.7%) did not show a preference during the first 3 min of the experiment; Small and young: 71 vs 27 males, respectively, G-test p<0.001, G^2^=20.48, p<0.0001; 32 males (24.6%.) did not show a preference during the first 3 min of the experiment). A post hoc examination revealed that males’ preference of females with relative higher quality was resembles among all 4 dyads (2×2 Fisher exact test &Holm-Bonferroni correction, p>0.05 in all of the pairwise comparisons).

**Figure 5.**
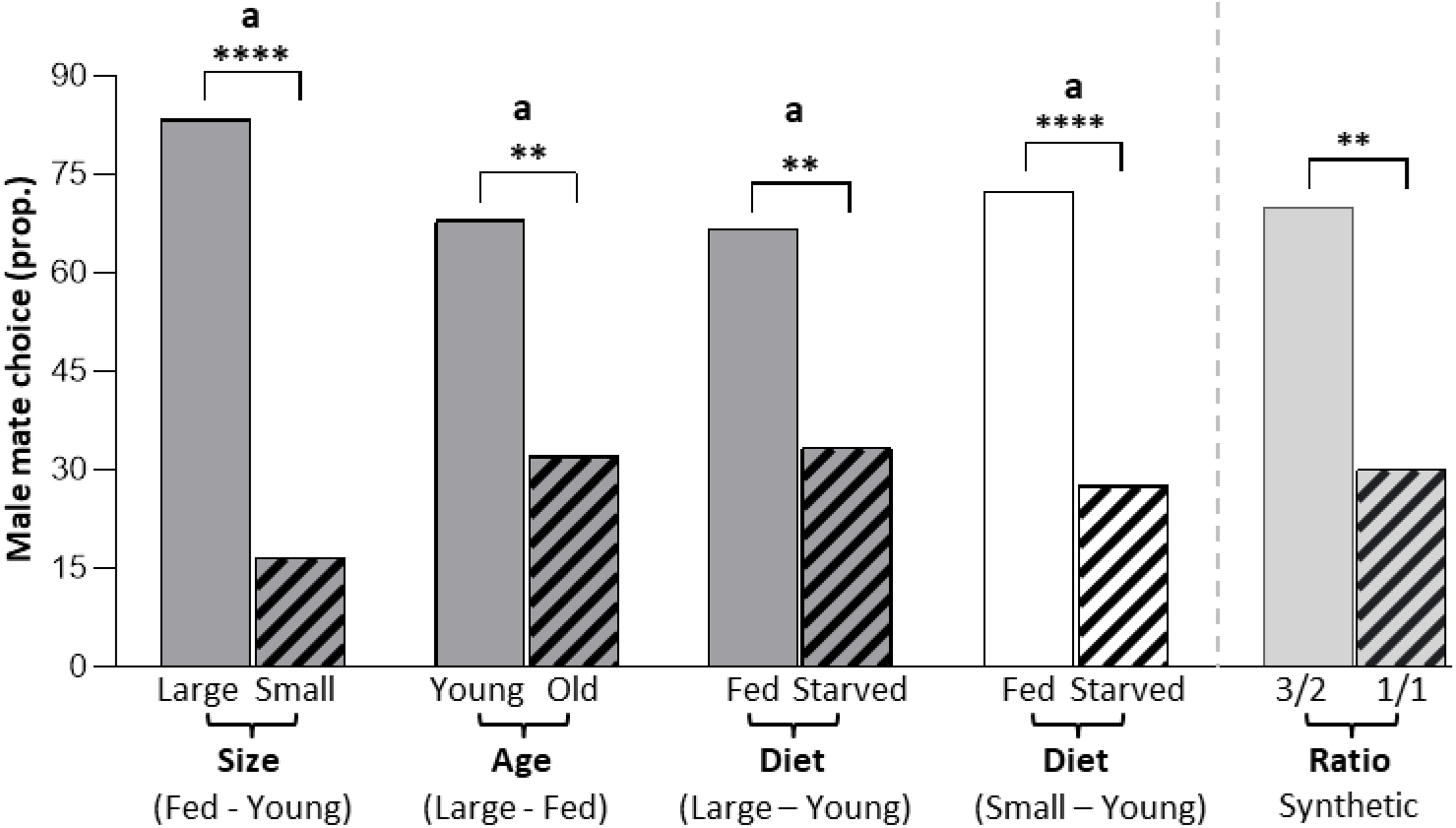
Number of pink bollworm males observed choosing the olfactometer arm through which the odor from the indicated groups of females flowed. (1) Within a group comparison, G^2^-test goodness of fit, asterisks indicate statistical significance of difference with ** p < 0.01, **** p < 0.0001. (2) Among groups comparison, Fisher exact test for multiple comparisons following by post-hoc comparisons using the Holm-Bonferroni method. Similar letters indicating for no significant differences.

Synthetic pheromone. Among the 66 males tested in the olfactometer, significantly more males chose the arm containing the pheromone blend of 0.6:0.4 ZZ:ZE than the arm having the 0.5:0.5 ZZ:ZE pheromone blend (42 vs 18 respectively) (G^2^=9.87, p=0.002) (Figure 5). Six males (9.1%.) did not show a preference during the first 3 min of the experiment.

## Discussion

A central request of the handicap principle in relation to sexual communication is that the sexual traits or its maintenance should be costly and thus positively correlated with the sender’s condition, allowing the receiver to base its mate choice on an honest, reliable signal (Grafen 1990; Johnstone 1995; Zahavi 1975). In accordance, several studies have provided evidence that male produced sex pheromones convey information to mate searching females, such as longevity (Johansson et al. 2005; Nieberding et al. 2012), satiation level (Foster and Johnson 2010), inbreeding status (Van Bergen et al. 2013; Westerman and Monteiro 2013); immunity (Rantala et al. 2003), and even sperm availability (Blaul and Ruther 2011). However, similar studies focusing on female produced pheromones and male mate choice are lagged.

In this study, in an attempt to close this gap we demonstrate that (1) female fecundity is correlated with their physical conditions such that female in good conditions, i.e. large, young or fed, have more offspring than females in bad conditions, i.e. small, old or starved. (2) The pheromone blends of the pink bollworm females accurately echo their current phenotypic conditions, and (3) males use this information when choosing a mate. Males preferred large females over smaller ones, young over old, large and fed females over large but starved, and small and fed females over small but starved. These mate preferences coincided with pheromone ratios that show more of the ZZ component in young, large and fed females and with the females’ reproductive potential. Age, size and satiation are physical characteristics that are typical for insect females in better physiological conditions and higher fecundity (Honěk 1993; Leahy and Andow 1994; Li et al. 2005; Wäckers et al. 2007).

Apparently, *P. gossypiella* females, in most conditions, produced similar amount of the full pheromone blend, except for females with the worst condition (small and starved), which produced significantly lower amount of pheromone. Using the pheromone characteristics in the gland of the pink bollworm females as a proxy for the characteristics of the pheromone emitted by calling females (Allison and Cardé 2016), we provided evidence that the information given by the amount of the pheromone alone, allows males to discriminate only among females in extremely poor conditions from other females. In contrast, more accurate information is provided through the pheromone ratio of components. Our results suggest a correlation between the female phenotypic conditions and the ZZ ratio in her pheromone, such that females in better conditions produce more of the ZZ component. Moreover, our finding suggests that the information conveys through both the pheromone amount and the ratio of components may more adequately inform the male of the female reproductive potential. This denotes a higher cost in producing more of the ZZ pheromone component, which only females in good physical condition can do.

Male preference for females in better conditions may have also obtained if females in good conditions put more efforts in increasing calling bouts (higher rate of the pheromone gland exposure per time unit) than females in bad conditions. Doing so, females in good conditions may assist flying males in targeting their pheromone patches. This phenomenon, if occurs, may indicate that calling is also a costly behavior. This may allow males to integrate the information emerged from the pheromone ratio with the female’s rhythm of calling to discriminate among females in good vs. bad conditions.

Difference in amount and/or ratio of the pheromone components in relation to fitness was found in previous studies of female choice of mates (Size - Chemnitz et al. 2015. Age - Nieberding et al. 2012. Diet - Blaul and Ruther 2011; Chemnitz et al. 2015; Fisher and Rosenthal 2006; Moraes et al. 2008). Examples for the cost of female produced pheromone are lagging behind that of males. Delisle and Vincent (2002) demonstrated that a female produced pheromone bears a cost since insecticide resistant female moths produced decreased amounts of sex pheromone. Barthel et al. (2015) showed a change in the ratio of the pheromone components in immune challenged females, and Foster and Johnson (2010, 2011) demonstrated the negative effect of low concentration of trehalose in the female hemolymph on pheromone production, while feeding on sucrose restored pheromone production. The effect of female size on characteristics of her pheromone has been detected in a few moth species (*Neoleucinodes elegantalis*, (Jaffe et al. 2007); *Lobesia botrana* (Harari et al. 2011), and age-dependent pheromone release has been demonstrated in various studies (Levi-Zada et al. 2020; Miller and Roelofs 1980; Noldus and Potting 1990; Raina et al. 1986).

Mate choice should be based on a signal that honestly reflects the signaler conditions that are relevant to the receiver; consequently, there is evidence that females choose their mates based on their pheromone characteristics. For example, older male burying beetles (*N. vespilloides)* invest more in parental care (Benowitz et al. 2013). In the butterfly *Bicyclus anynana* mid-aged males advertise good genes that allow for better survival on the one hand (Manning 1985) and assure no sperm depletion on the other. In some lepidopteran species, males advertise the quality of their nuptial gifts via their pheromone (Dussourd et al. 1991; Eisner and Meinwald 1995).

Similarly, Size (Harari et al. 2003; Honěk 1993; Li et al. 2005; Xu and Wang 2009), age (Moore and Moore 2001; Torres-Vila et al. 2002; and in spiders Waner et al. 2018) and diet (Foster and Johnson 2010; Wäckers et al. 2007) have large impacts on females’ fecundity on the one hand and significantly affect pheromone characteristics on the other (Harari et al. 2011). Therefore, by perceiving the present reproductive potential of a female, via her currently released pheromone, males should increase their direct reproductive success, when exert a choice of females (Biaggio et al. 2016; Waner et al. 2018), as was found in this study.

Interestingly, the pheromone amount in the glands did not fluctuate much among females in most conditions. This finding differs from various studies suggesting that stress affects the amount of pheromone produced by females (Delisle and Vincent 2002; Foster and Johnson 2010; Foster and Johnson 2011). In their model, Umbers et al. (2015) predicted that old virgin female moths should increase their signaling efforts by expending signaling time but not pheromone titer. This is based on the vast majority of studies reporting that old females did not increase their pheromone amount in the gland, despite the risk of remaining unmated. Although the relation between the pheromone amount in females’ gland and their calling period has not much investigated (Umbers et al. 2015), the notion that old females do not increase pheromone production, regardless of the risk to remain unmated (e.g. Gemeno and Haynes 2000; Levi-Zada et al. 2020), suggests that pheromone production bears a high condition-dependent cost. Alternatively, larger pheromone amount may benefit the signaling female as it transfers for longer distance and may attract more males (Greenfield 1981). However, the amount, per-se, may not serve as a honest indication for female quality as its concentration changes in relation to the distance in time and space of its source (Valeur and Löfstedt 1996; Willis and Arbas 1991), whereas the ratio of components remain constant for long distances along the pheromone plume (Kaissling 1996; Linn et al. 1987; Meng et al. 1989).

In the present result, females in most physical conditions produced similar amounts of pheromone, which significantly differed from that of females in harsh conditions (small and starved). This is in accordance with the handicap principle claiming an unequal cost of advertising, such that for a similar signal intensity higher cost is inflected on individuals in poor conditions than on individual in good conditions (Zahavi 1975, 1977; Grafen 1990; Johnstone 1995). Here, the failure of small and starved females to produce the pheromone in large quantities suggests that the production of the pheromone, in itself, is costly; whereas females under most experimental conditions were able to tolerate the cost, females under extreme stress could not. Presumably, a higher cost, and thus, a better signal for quality, involved in producing more of the ZZ pheromone component, which only females under ideal conditions can afford. Currently the cost of diverting the pheromone ratio away of the “population mean” is not fully understood, but it has been suggested that an increase in a specific enzyme’s levels that is involved in the pheromone biosynthesis, affects the ratio of the pheromone components (Blomquist et al. 2012). Accordingly, Dou et al. (2019) found different expressed genes of the pink bollworm that are associated with females having high ZZ ratios in their pheromone.

Interestingly, whereas the GC failed to decipher between pheromone characteristics of fed and starved large females, males significantly preferred large and fed females over starved female. Doing so, they gained females with significantly higher reproductive potential as was found in the current study.

The results of this study join the rapidly accumulating evidence that the population variance in pheromone characteristics is subjected to directional sexual selection, whereby male choosiness promotes the increase of certain components of the female sex pheromone (*Lobeisa botrana*, Harari et al. 2011). It is of interest to test whether pheromone signal quality is not unique to the species studied here, but is a general trait of species in which females release sex pheromones and males bear the cost of searching for a mate and mating.

## Author Contributions

A. Gonzalez-Karlsson and Y. Golov sharing first authorship. Conceptualization: A. Harari, A. Liberzon, R. Gurka, R. Jurenka, V. Soroker and Y. Golov. Data curation: Gonzalez-Karlsson, Y. Golov, A. Moncaz, E. Halon. Funding acquisition: A. Harari, R. Jurenka, A. Liberzon. Statistical analyses: Y. Golov. Methodology: Y. Golov, A,. Harari, A, Liberzon, R. Gurka. Supervision: L. Liberzon, R. Gurka, A, Harari. Writing – original draft: Y. Golov and A. Gonzalez-Karlsson. Writing – review &editing: A. Liberzon, R. Gurka, R. Jurenka., H. Steinitz, R. Horowitz, I. Goldenberg, V. Soroker. A. Harari.

## Acknowledgement

We thank Michal Axelrod, Ariela Niv and Nikolay Meltser for their valuable field assistance and inspiring discussions. This study is supported by the U.S.- Israel Binational Science Foundation (BSF) research grant award 2013399, to AH, AL and RG, and the United States-Israel Binational Agricultural Research Development fund (BARD) research grant award IS-4722-14 to AH, VS, and RJ.

## Appendixes

In attempt was made to define the female quality by including her physical condition (i.e. her size, age and diet), her fecundity, and her pheromone characteristics (i.e. amount and ratio of components) into one analysis. A multivariate analysis was performed, using the descriptive technique of multiple correspondence analysis (MCA). For MCA analysis two steps are required: (1) A homogeneous multiway table, and (2) data pooling. (1) in order to meet the requirement of homogeneous multiway table, the size of the data sets of both types of parameters, (i) female fecundity (number of eggs) and (ii) pheromone traits (amount and ZZ/ZE ratio) was equalized, in each of the eight groups (see Appendix. 1). That is, for each the females’ physical condition (large, young and fed), the sample-size was equalized using a simple random sampling (without replacement) on the limiting factor (data with smaller sample size). (2) Due to the methodological limitation of the GC analysis (comparing within dyads only), the MCA did not explore the associations among the dyads. Therefore, following the four dyads (see Table. 2), the number of oviposited eggs was pooled separately for each treatment, either size (large and small), age (young and old) diet (fed and starved) and small and diet (small fed and small starved). Then, for each female, we calculated her proportional contribution to the egg pool (the total number of eggs of all females in her dyad). In addition, we defined Boolean variables (marked as ‘+’ and ‘-’) relative to each of the physical conditions: (+) to the better physical condition in the dyad, and (-) to the relatively inferior physical condition in the dyad.

**Appendix 1.** Data processing for the MCA.

| Treatment | Size |  |  |  | Age |  |  |  | Diet |  |  |  | Diet |  |  |  |
| --- | --- | --- | --- | --- | --- | --- | --- | --- | --- | --- | --- | --- | --- | --- | --- | --- |
| Shared features | Young & Fed |  |  |  | Large & Fed |  |  |  | Large & Young |  |  |  | Small & Young |  |  |  |
| Polled females | Large & Small |  |  |  | Young & Old |  |  |  | Fed & Starved |  |  |  | Fed & Starved |  |  |  |
| Sub-groups | Large |  | Small |  | Young |  | Old |  | Fed |  | Starved |  | Fed |  | Starved |  |
| Adjusted feature | Eggs # |  | Eggs # |  | None |  | Eggs # |  | Eggs # |  | Eggs # |  | Eggs # |  | Eggs # |  |
| Comparison | O | P | O | P | O | P | O | P | O | P | O | P | O | P | O | P |
| N | 31 | 13 | 30 | 10 | 27 | 27 | 31 | 28 | 17 | 13 | 29 | 16 | 30 | 24 | 28 | 11 |
| Total counts | 23 |  |  |  | 55 |  |  |  | 29 |  |  |  | 35 |  |  |  |

**Appendix 2.**
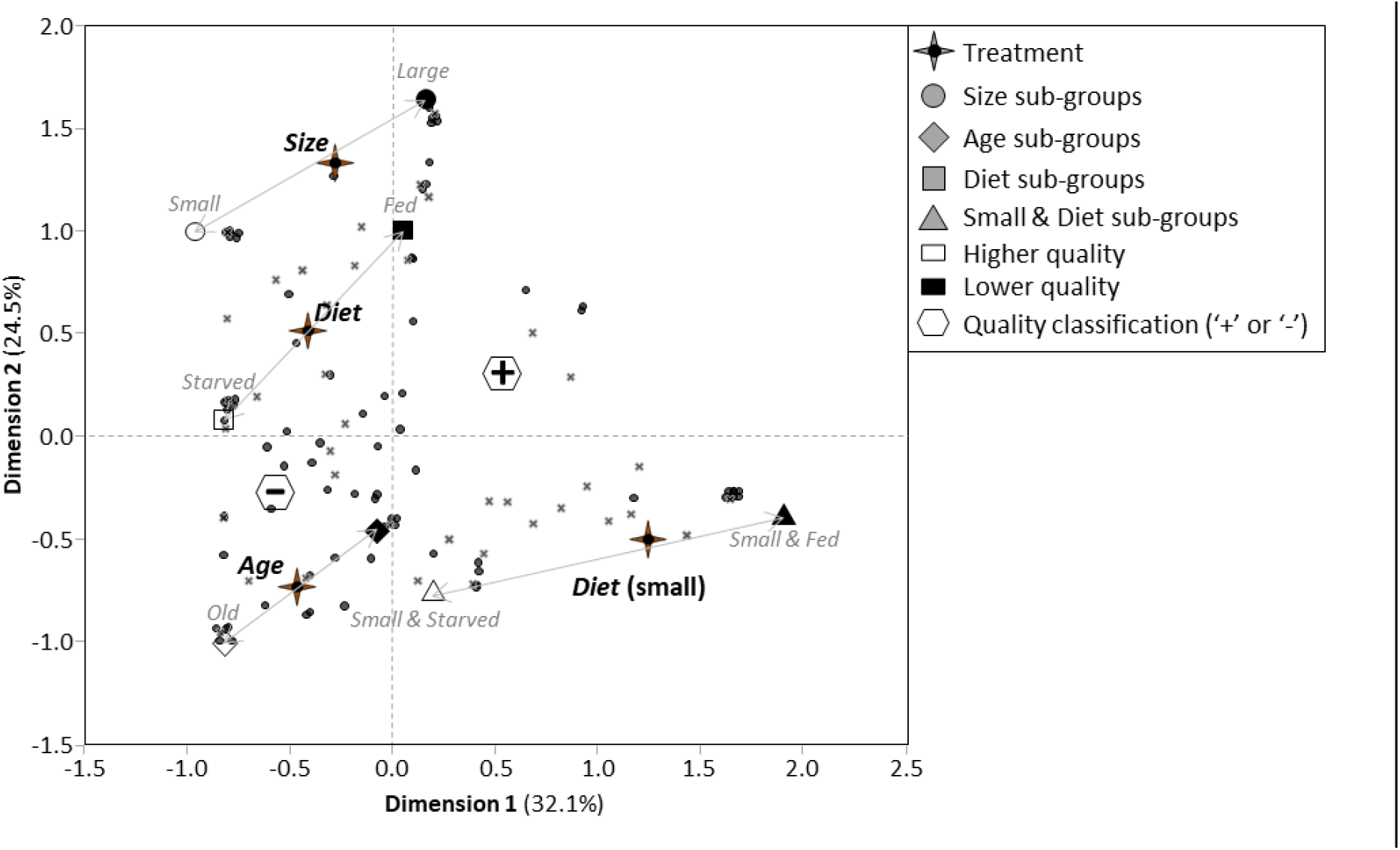
Association between female’s physical condition and her tested traits. Comparability is displayed in a 2-D Euclidean space. Independent variables (x): x1=‘size’, x2=‘age’, x3=‘diet’, x4=‘diet of small’ (four groups), supplementary variables: females of higher (‘+’) and lower (‘-’) quality in each of the four dyads. Dependent variables: y1=eggs number, z1= pheromone amount and z2= proportional pheromone ZZ/ZE ratio. Multivariate technique of multiple correspondence analysis (MCA).

